# Environmental, social, and morphological drivers of fission-fusion dynamics in a social ungulate

**DOI:** 10.1101/2022.09.22.508899

**Authors:** Morgane Le Goff, Jack G. Hendrix, Quinn M. R. Webber, Alec L. Robitaille, Eric Vander Wal

**Affiliations:** Behavioural ecology and wildlife management, Université de Bourgogne, Dijon, France; Cognitive and Behavioural Ecology Interdisciplinary Program, Memorial University of Newfoundland, St. John’s, NL, Canada; Department of Integrative Biology, University of Guelph, Guelph, ON, Canada; Department of Biology, Memorial University of Newfoundland, St. John’s, NL, Canada

**Keywords:** Fission-fusion, *Rangifer tarandus*, body size, predation, landscape heterogeneity, sociality

## Abstract

Social groups exist because individuals within the group accrue a net benefit from sharing space. The profitability of sociality, however, varies with ecological context. As ecological context varies, tension emerges among the costs and benefits of social grouping. Fission-fusion societies are fluid in their group dynamics across spatial and temporal contexts, permitting insights into how context affects whether animals choose to join or depart a group. We tested four non-mutually exclusive hypotheses driving variation in fission and fusion in caribou: the risky places, environment heterogeneity, activity budget, and social familiarity hypotheses. The risky places hypothesis predicts animals are unlikely to diffuse when habitats are open and risk of predation is elevated. The habitat heterogeneity hypothesis predicts that fission is more likely in a heterogeneous landscape due to the rising conflicts of interest between group members. The activity budget hypothesis predicts dyads associate by body size due to similar food passage rates. The social cohesion hypothesis predicts that familiar individuals are less likely to fission. We tested the hypotheses using time-to-event (time before fission) analyses and a linear model that assesses spatial, social, and body size relationships among female caribou (n = 22) on Fogo Island, Newfoundland, Canada. Contrary to our prediction for risky places, probability of fission was not influenced by habitat openness. The hypothesis of environmental heterogeneity was partially supported, as caribou remained less cohesive in environments with a higher richness of habitats. No direct evidence emerged to support the activity budget hypothesis. However, it appears that caribou maintain the strongest social bonds among variably sized individuals and these social bonds do decrease the propensity to split. Collectively, our findings showed that social interactions may depend not only on individual identity and characteristics, but also the spatial context in which these interactions occur.

## Introduction

Changing ecological contexts influence the costs and benefits of animal social behaviours (Webber & Vander Wal, 2018). For gregarious species that experience rapid or ongoing changes in ecological contexts, social groups can range from stable with limited inter-group movement to dynamic fission-fusion societies with frequent merging and splitting (Aureli et al., 2008). Animal groups are predicted to reach an optimal size that maximizes fitness within a given context (Carter et al., 2009; Webber & Vander Wal, 2018; Webber & Vander Wal, 2021). For example, risky habitat constitutes a key ecological context that can result in group fusion to dilute predation risk (Moll et al., 2016). Alternately, complex habitats provide cover from predators, can result in predator confusion, and are thus predicted to result in group fission (Fortin et al., 2008). Furthermore, within social groups, conflicts can also arise between individuals, affecting fission (Conradt & Roper, 2000). For example, according to their body size, some ungulate species allocate time differently to foraging based on size-specific digestion efficiency (Ruckstuhl, 2007). As a result, there is a mismatch between group members in the time required for foraging and digestion (Ruckstuhl & Neuhaus, 2002). Consequently, variation in intrinsic requirements of individuals in the group drives fission into subgroups (Conradt & Roper, 2000). Moreover, pre-existing social relationships may also affect fission. For example, familiarity between individuals may minimize fission of social groups (Carter et al., 2013). Here, we consider the variation in the ecological (i.e., perceived predation risk and habitat heterogeneity), morphological (i.e., body size), and social (i.e., familiarity) contexts of a gregarious ungulate and the implications of these contexts on fission-fusion dynamics.

Predation risk related to habitat openness influences group size of prey species and drives fission-fusion dynamics (Fortin et al., 2009). Group living offers anti-predator benefits (Krause et al., 2002) such as higher detection of predators (Leuthold, 1977) and predator harassment (Berger, 1979). Foraging animals aggregate in groups and use collective defenses in risky habitats to mitigate predation risk (Molvar & Bowyer, 1994). The risky places hypothesis suggests that such anti-predator behaviour differs based on the long-term background risk associated with different environments, irrespective of short-term pulses of risk or safety (Moll et al., 2017). Indeed, predation risk is often associated with habitat openness, as it visually exposes prey to predators (Ebensperger & Wallem, 2002; Mao et al., 2005). As such, groups of prey can have different strategies to reduce predation risk. In some species, individuals may forage in large groups in areas where food is more profitable, but the risk of being predated is high, i.e., open habitat. Meanwhile, individuals of other species may forage in smaller groups in safer areas where the food is less profitable, but the risk of predation is lower, i.e. in closed habitat or next to cover (Lima & Dill, 1990). For example, spider monkeys (*Ateles fusciceps*) fuse into larger groups when occupying open habitats perceived to be high-risk, e.g., mineral licks (Link & Di Fiore, 2013). Under high predation risk, large groups also tend to have higher overall rates of vigilance so that on a per capita basis individuals spend more time feeding while reducing the group-level predation risk (Lima, 1995). Animals therefore adopt a range of behavioural strategies to reduce the perceived risk of predation through space and time (Gaynor et al., 2019).

Landscape heterogeneity also affects decision-making, group movement, and variation in predation risk. An uneven distribution of resources and predators increases the potential for a conflict of interest within a group (Sueur et al., 2011). For example, conflict of interest can arise from a preference in a foraging direction, e.g., move toward food patch A or B. In this case, fission into two groups is likely, since the average direction between A and B will not profit either sub-group (Sueur et al., 2011). Individuals that are unable to synchronize their activities (e.g., foraging, travelling, resting) are predicted to fission into separate groups (Ruckstuhl & Neuhaus, 2002). The environmental heterogeneity hypothesis predicts a higher probability of fission in heterogeneous environments due to a broader range of options for the different needs and motivations of individuals in groups. Winnie et al. (2008) found that heterogeneity in quality and quantity of forage explained fission-fusion dynamics in buffalo (*Syncerus caffer*). In addition to the external drivers of fission-fusion such as predation pressure and habitat heterogeneity, intrinsic traits can play a role in fission-fusion dynamics.

The activity budget hypothesis has specific predictions for ungulates where variation in body size affects synchronization of behaviour. Body size is an important intrinsic trait that generates conflict among ruminant group members and alters group cohesion. For an individual to synchronize their activities with other group members, they may have to compromise their own activity budget, which can be costly in groups that include members of different age, sex, or body size (Bon et al., 2006). The allocation of time to different activities is more likely to vary between individuals with different nutritional requirements. In particular, individuals of different body sizes can have varying digestion efficiency in ruminants, which could result in subgroups of similar-sized individuals (Bon et al., 2006; Ruckstuhl, 2007). In sexually dimorphic ungulates, smaller individuals are less efficient at digesting fibrous food and as a result, smaller individuals forage for longer and more selectively than larger individuals. This results in a segregation of individuals where some spend more sedentary time ruminating (Ruckstuhl & Neuhaus, 2002). According to the activity budget hypothesis, differences in activity budgets could explain sexual segregation in size-dimorphic ungulates (Ruckstuhl & Neuhaus, 2002; Bon et al., 2006).

Although not as common, the tendency to synchronize activities by size can also occur within groups of males or females. For example, pairs of female Gasconne beef cows (*Bos taurus*) of similar weight, and thus size, were more synchronized than pairs of dissimilarly sized females (Šárová, Špinka, & Panamá, 2007). Among male bighorn sheep (*Ovis canadensis*), groups composed of similar-sized individuals are more synchronous than groups composed of individuals of varying sizes, presumably because individuals of different sizes must pay a metabolic cost if they want to stay in cohesive groups (Ruckstuhl, 1999). Assortment by size allows individuals of similar needs to stay cohesive, without having to pay the cost of synchrony, which can impair foraging efficiency (Meldrum & Ruckstuhl, 2009; Aivaz & Ruckstuhl, 2011).

Another factor likely to affect fission and fusion is social familiarity among individuals. Social familiarity occurs when two individuals engage in affiliative interactions, e.g., spending time together, huddling, cooperatively foraging, considerably more often and over greater periods than other individuals (Brent et al., 2014). For example, social familiarity influences fission-fusion dynamics in giraffes (*Giraffa camelopardalis*), where adult females giraffe spend more time with preferred individuals (Malyjurkova et al., 2014), that are not necessarily kin (Carter et al., 2013). Over longer periods of time, close associations facilitate social learning of foraging tasks (Benskin et al., 2002; Figueroa et al., 2013) or anti-predator behaviours (Kavaliers, Colwell, & Choleris, 2005). The use of social information can therefore be an asset in heterogeneous landscapes, which are increasing in frequency as anthropogenic disturbances are generating fragmentation of natural landscapes; social information is thus particularly beneficial in these areas (Fletcher & Sieving, 2010).

Woodland caribou (*Rangifer tarandus*), hereafter caribou, are gregarious ungulates that live in loose fission-fusion societies (Lesmerises, Johnson, & St-Laurent, 2018) and form groups whose abundance (Edmonds, 1998) and strength of social associations vary seasonally, i.e., smaller groups in summer and larger groups in winter (Robitaille et al., 2021; Webber & Vander Wal, 2021). Female caribou forage in larger groups in risky habitats and increase their vigilance compared to safer habitats (Bøving & Post, 1997). Moreover, caribou tend to select habitats that reduce their predation risk (Basille et al., 2015; Bastille-Rousseau et al., 2016), especially during calving (Bonar et al., 2020).

Our objective was to determine how predation risk, environmental heterogeneity, body size, and social familiarity among female caribou affect fission-fusion dynamics. We tested four hypotheses, which beget four non-mutually exclusive predictions of fission events (Figure 1):

**Figure 1:**
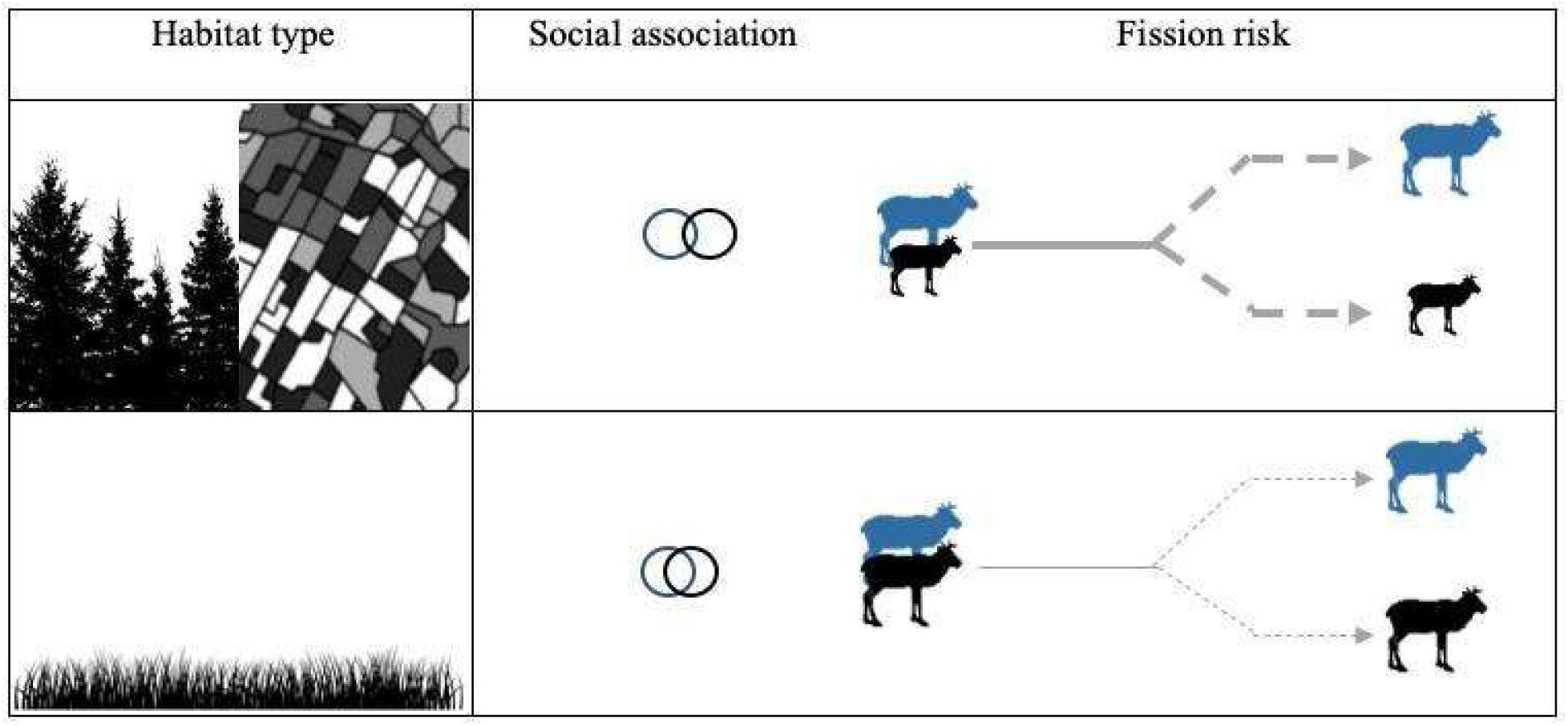
Schema of the predictions associated with fission probability tested in our study. Different habitat types are represented by a forest (closed habitat), a mosaic of habitats (heterogeneous landscape) and a meadow (open habitat). Solid lines represent dyad steps and dashed lines individual paths taken after fission. Lines are thicker with increasing fission risk. The degree of attachment of circles refers to the degree of association between caribou. The more circles overlap, the stronger the social association.

1. The risky places hypothesis suggests that groups of prey fuse in risky habitats and split in safer habitats as an anti-predator strategy associated with variation in the inherent risks of different environments (Moll et al., 2017). Therefore, we predicted that caribou groups would split less in open habitats where predation risk is assumed to be higher (P1).
2. The environmental heterogeneity hypothesis predicts that complex environments make it more difficult to remain in cohesive groups because members of a social group have different foraging needs and requirements, which can lead to conflicts in decision-making (Fortin et al., 2008). We therefore predict that groups of caribou will be more likely to split in heterogeneous environments (P2).
3. The activity budget hypothesis predicts that individuals with similar energetic needs, and therefore similar body sizes, form cohesive groups and separate from animals with different needs because synchronizing their behaviour can be costly (Ruckstuhl, 1999). Thus, we predict that individuals that are more similar in their body size will be more likely to stay fused longer than individuals that are more dissimilar in their body size (P3).
4. The social familiarity hypothesis predicts the probability of fission decreases for dyads with stronger social familiarity because remaining together can provide them with a fitness benefit (Brent et al., 2014). We predict that dyads with higher pairwise simple ratio index (SRI), a metric of social association, will be less likely to split than dyads of less familiar individuals (P4).

In addition to our models of fission events, we also tested how space use and home range overlap may influence social associations. By definition, animals that aggregate together must share at least some of the same home range, and there is no opportunity for fission events to occur if animals have not already fused, i.e. coexistence is space is a prerequisite for these social associations. Moreover, based on the activity budget hypothesis, familiarity ought to be explained by similarity in size, with greater familiarity between individuals of similar size. We thus predicted that SRI between caribou pairs will increase with greater home range overlap, and decrease as the difference in body sizes increases (P5).

## Methods

### Study area and subjects

We studied the social behaviour of caribou on Fogo Island, located off the Northeastern coast of Newfoundland, Canada (Latitude: 49° 39’29.39” N; Longitude: 54° 10’7.80” W). Caribou were introduced to Fogo Island in the 1960s as part of a series of introductions throughout Newfoundland (Bergerud & Mercer, 1989) and the population currently consists of ∼300 individuals (Newfoundland and Labrador Wildlife Division, *unpublished data*). Although caribou are predated by black bears (*Ursus americanus*) and coyotes (*Canis latrans*) on the island of Newfoundland (Bastille-Rousseau et al., 2016), only coyotes are present on Fogo Island (Huang et al., 2021). Caribou in Newfoundland generally favour open habitats (Bergerud, 1974) for their abundant forage and avoid forested habitats that are difficult to access and offer few forage opportunities (Fortin et al., 2008). Caribou diet changes seasonally based on the accessibility of resources. During summer, caribou are generalists, foraging on shrubs, lichens, sedges, and herbaceous plants (Bergerud & Nolan, 1970; Webber et al., 2022). During winter, they either dig holes in the snow termed craters and consume terrestrial lichens, or forage on arboreal lichens when access to terrestrial lichens is hindered by the snow depth or its hardness (Johnson, Parker, & Heard, 2001). We focused our study on winter (2017-2019), defined as 1 January to 16 March, which corresponds to previous models of caribou social behaviour, movement, and habitat selection (Bastille-Rousseau et al., 2016; Peignier et al., 2019).

### Caribou capture and collar data

Newfoundland and Labrador Wildlife Division carried out the capture of adult female caribou (*n* = 31) between 26 March and 20 April 2016-2018 using the immobilizing agent Carfentanil, administered via dart gun. All animal captures and handling procedures were consistent with the American Society of Mammologists guidelines and were approved by Memorial University Animal Use Protocol No. 20152067. Caribou were fitted with global positioning system (GPS) collars (Lotek Wireless Inc., Newmarket, ON, Canada, GPS4400M collars, 1.250 kg), which collected location fixes every two hours. Of the original 31 caribou, 9 were removed from subsequent analyses either due to collar failure or death. Prior to analyses, we removed the erroneous GPS fixes resulting from malfunctioning collars following the screening method of Bjørneraas et al. (2010). This method relies on previous knowledge of the study species and excludes implausible fixes like those further than a predefined maximum distance an animal could travel, and fixes representing spikes in the movement trajectory. We assumed the sample of collared females was random among adult females and the measures of social familiarity (see below) between caribou were an unbiased representation of associations in the broader population. Overall, we used the locations of 11 caribou in 2017, 16 in 2018 and 13 in 2019.

Body measurements were recorded upon capture. Specifically, we measured total length from the end of the upper lip to the last vertebra of the tail, heart girth as the circumference behind the forelegs, and neck girth as the circumference where the GPS collar is fitted. Heart girth and total body length are common measurements used to assess body size in ungulates (McElligott et al., 2001; Cook, Cook, & Irwin, 2003). Body size along with body condition are two components of body mass. Heavier individuals are typically larger than lighter individuals and among similar-sized individuals, heavy individuals have better body condition (Toïgo et al., 2006). In our study we did not have access to body mass or body condition data, so we used body size as a proxy for body condition and weight. For subsequent analyses, we used the total body length (range: 174–216 cm) as a proxy for body size instead of heart girth (range: 110– 131cm) because total body length was more variable. For individuals with multiple measurements of body length, we calculated the average length for subsequent analyses. All the statistical analyses were performed with R (R Core Team, 2021).

### Calculating dyads

We used the *spatsoc* package (Robitaille, Webber, & Vander Wal, 2019) to group GPS locations in time (within 5 minutes) to account for temporal variation between GPS fixes of different animals in the same time step. We defined dyads as times when individuals were located within a 50m buffer of one another for at least two relocations, following Lesmerises et al. (2018). The same individuals could therefore be a part of different dyads at different times. We used the median location of the two individual’s GPS fixes as the dyad location at a given time, to calculate subsequent landscape measures (see below) and as a unit for subsequent analyses.

To delineate fission and fusion events, following Lesmerises et al. (2018), we used the dyadic centroid to represent the combined dyadic step. We first defined fusion as events where two individuals were within 50m for at least two consecutive time steps. We then defined fission as events where individuals previously in a dyad were more than 50 meters apart for at least two consecutive time steps. In cases where dyads were together before and after one missing GPS relocation (from one individual in the dyad), we assumed the dyad remained together (see Figure S1). Our analyses primarily focused on fission events, whereas fusion events were not the explicit response variable in any of our models.

We described the strength of association between two caribou through years using the simple ratio index (Cairns & Schwager, 1987): where *x* is the number of times individuals A and B were within the 50 meters threshold and *yAB* is the number of simultaneous fixes from individuals A and B that were separated by more than 50 meters (Farine & Whitehead, 2015). Higher values of SRI reflect stronger associations, and thus social familiarity, between individuals.

### Home range area and overlap

We estimated each individual’s home range in each year using 95% kernel density estimates from the *adehabitatHR* package (Calenge, 2006). To calculate home range overlap, we extracted each individuals’ kernel and calculated the utilization distribution (i.e. probability distribution defining the animal’s space use) overlap index (UDOI) between dyads to quantify overlap in terms of space-use sharing (Fieberg & Kochanny, 2005). UDOI values in our analyses ranged from 0 (no overlap) to 1.46 (high degree of overlap).

### Habitat and land cover classification

The land cover data of Fogo Island consisted of nine habitat types at 30 m spatial resolution (Integrated Informatics Inc., 2014). Habitats included wetland, broadleaf forest, conifer forest, conifer scrub, mixed wood forest, rocky barrens, water/ice, lichen barrens, and anthropogenic areas. We used all nine habitats types for the subsequent calculations of heterogeneity metrics (P2), but we grouped habitats into two categories for our analysis of predation risk (P1): closed habitat (broadleaf forest, conifer forest, conifer scrub, and mixed wood forests), and open habitat (wetland, water/ice, rocky barrens, lichen barrens, and anthropogenic areas). We used habitat openness as a proxy for perceived predation risk with open habitat representing riskier areas than forested ones. The proportion of open habitat was calculated at the beginning of each dyad step in a 200 m buffer around the centroid of the locations of the dyad.

To account for habitat heterogeneity, we described two aspects of a landscape: spatial configuration and spatial composition (Li & Reynolds, 1993). We calculated the contagion index, which is an aggregation metric to describe habitat configuration, the arrangement of the different land cover types. We also calculated the Shannon index to describe habitat composition. The contagion index is a measure of spatial distribution and intermixing of patches, which describes the probability that two randomly chosen adjacent pixels belong to two different habitat classes. Hence, it can be perceived as a measure of habitat fragmentation (Ricotta, Corona, & Marchetti, 2003). The contagion index (McGarigal et al., 2002) is calculated as:

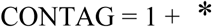

with *p*_*q*_ the adjacency table (i.e., matrix showing the frequency with which different pairs of habitat class appear side-by-side on the map) for all habitat classes divided by the sum of that table and *t* is the number of habitat classes in the landscape. Values range between 0 and 1 with values close to 1 associated with homogeneous landscape, with few large contiguous patches of the same habitat class, whereas values close to 0 characterize heterogeneous landscapes with many small patches, highly dispersed (McGarigal et al., 2002).

The Shannon index (Shannon, 1948) is a common measure of habitat diversity that accounts for both abundance and evenness of habitats and is calculated as:

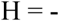

where *S* is the number of habitat classes and *p*_*i*_ the proportion of pixels belonging to the *i*th cover class. Diversity increases with increasing values of H. We computed both the contagion and Shannon’s indices within a 200m buffer around the centroid of each caribou dyad location.

### Statistical analyses

To assess our predictions, we conducted two separate model sets. First, we modelled the probability of dyad fission based on habitat openness (P1), habitat heterogeneity (P2), difference in body size (P3) and social association (P4). Specifically, we used a time-dependent Cox proportional hazards model using the package *coxme* that account for mixed effects (Therneau, 2020). In our Cox proportional hazards model, each time interval was represented by a time step for a dyad and the covariates included, the proportion of open habitat within a 200m buffer of the dyad, the Shannon index and the contagion index within a 200m buffer of the dyad, the difference in body size and the dyadic SRI. For each time step, the status (i.e., survival) of a dyad was assessed, i.e., either together or split. Landscape metrics were specific to each unique dyad step, whereas the SRI for dyads was constant through time within each year. Since the same dyad could be associated at different occasions throughout the three years of study, we included dyad ID and year as random effects.

Second, we modelled pairwise association strength as a function of home range overlap and similarities in body size (P5) to test whether female caribou preferentially associate with similar-sized individuals. Data for this model set was based on aggregate annual measures of association (i.e., SRI), body size, and home range overlap. Specifically, we used a linear mixed model using *lme4* with pairwise SRI as the response variable, the difference in body size and home range overlap between dyads as fixed effects, the interaction of body size difference and home range overlap, and dyad ID and year as random effects (Bates et al., 2015). We removed dyads with no home range overlap because these individuals did not have an opportunity to associate and therefore no home range overlap automatically results in a shared SRI of zero. We square-root transformed SRI to improve the requirements of normality and homoscedasticity.

For all analyses, we used the Akaike information criterion (AIC) to select the most parsimonious model (Akaike, 1981) and set the threshold for significant effects to p < 0.05.

## Results

The Fogo Island caribou population displayed characteristics of a fission-fusion system, with fusion events lasting from hours to weeks (median = 6 h; range = 4 h - 17.6 days). We recorded 1617 fission-fusion events during the study period, with an average of 549 ± 137 (range = 457– 705) events per winter. In total, 93% of fission events occurred in open habitats, while 7% of fission events occurred in closed habitats. On average, 56 ± 26 (range = 40–85) unique dyads per year were formed.

The Cox proportional hazards model highlighted potential environmental and social factors that influence fission. Of the models considered, the most supported using AIC model selection included social familiarity (i.e., the SRI), difference in body size between individuals in the dyad, Shannon index, contagion index, and habitat openness (for model selection results see Table S1). There were several other highly ranked candidate models (ΔAIC < 3), all of which comprised the same fixed effects as our most supported model while including additional interaction terms between these predictors. None of the interaction terms had significant effects, and the main effect estimates of our top model were unchanged until those variables were included in an interaction, at which time the effect disappeared. We have thus chosen to focus our interpretation on this top model, as these additional interactions do not provide any additional insight in our analyses.

The probability of fission increased with increasing Shannon index but was not influenced by habitat openness, contagion index, or difference in body size (Table 1). The probability of fission decreased with higher dyad SRI (Table 1). Together, these results suggest that caribou were more likely to fission in landscapes with various land cover types regardless of their configuration, while dyads stayed together for longer when they were more familiar with one another.

**Table 1:** Results from the most parsimonious Cox proportional hazards model with hazard ratios (HR) and their 95% confidence interval (CI) explaining the fission probability of dyads between 2017 and 2019 (n = 1617). HR >1 implies an increasing risk of fission, while HR <1 implies a lesser risk of fission. If the CI includes 1, then the HR is not significant. Significant results are presented in bold. Model selection results are presented in Table S1.

| Variable | $\beta$ | SE | HR | 95% CI | p-value |
| --- | --- | --- | --- | --- | --- |
| Habitat openness | 0.400 | 0.217 | 1.495 | [0.976 - 2.283] | 0.06 |
| Difference in body size | 0.004 | 0.007 | 1.004 | [0.99 - 1.019] | 0.58 |
| **Shannon Index** | **0.517** | **0.222** | **1.677** | **[1.085-2.591]** | **0.002** |
| Contagion Index | 0.401 | 0.426 | 1.493 | [0.648-3.444] | 0.35 |
| **SRI** | **-1.703** | **0.639** | **0.182** | **[0.052-0.637]** | **0.01** |

In our linear mixed model of social association strength, the difference in body size in a dyad of caribou and their home range overlap explained their shared dyad SRI (Table S2). The interaction between difference in body size and home range overlap suggests that caribou that shared a larger portion of their home range were more closely associated when they had a greater difference in body size (LMM; *p* < 0.01; *z* = 2.85; *β* ± se = 0.007 ± 0.003; Rm^2^ = 0.61; Figure 2, Table S2).

**Figure 2:**
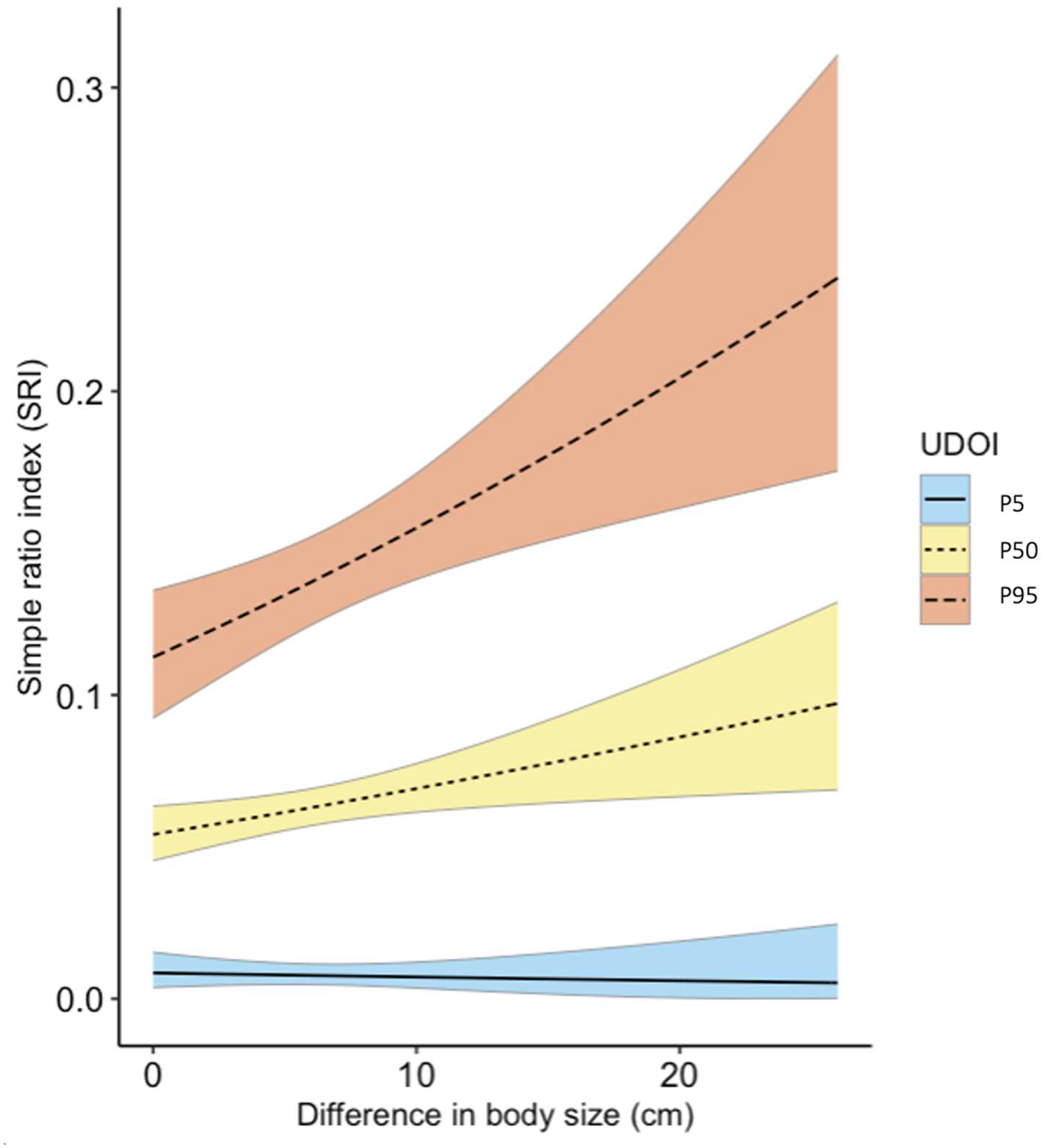
Changes in simple ratio index (SRI), measuring strength of social association, as a function of home range overlap (Utilization Distribution Overlap Index, UDOI) and difference in body size (cm) in caribou dyads, following the linear mixed model. UDOI was analysed as a continuous measure in the linear model, but is split into three values here for graphical purposes. Different colors represent the 5_th_ (blue), 50_th_ (yellow) and 95_th_ (blue) percentiles of UDOI to better visualize the change in SRI with its associated explanatory variables. Shading around each solid line is 95% confidence interval.

## Discussion

Factors driving fission-fusion dynamics are related to the social and ecological environments (Sueur et al., 2011). We tested the effects of habitat openness as a proxy for perceived predation risk, landscape heterogeneity, social familiarity among individuals, and similarity of body size on fission-fusion dynamics in caribou. In contrast to predictions from the risky places hypothesis, the probability of dyad fission was not greater in open habitats. We found no direct support for the activity budget hypothesis. Body size did not influence the risk of fission.

However, dissimilar body size and home range overlap collectively explained the strength of social association. Risk of fission decreased with increasing social association and increased in more heterogeneous landscapes.

Based on the risky places hypothesis, we predicted predation risk to drive fission-fusion dynamics by promoting fission in closed habitats (P1). Contrary to our prediction, the probability of fission was similar in open and closed habitats. Habitat openness influences group size for caribou such that larger groups tend to form in more open habitats (Webber & Vander Wal, 2021). While groups may indeed be larger in open habitats, the probability of fission is not associated with habitat openness. A potential explanation is that more open habitats facilitate groups to remain fused to exchange information about foraging sites (Peignier et al. 2019) and maintain high predator vigilance (Lima, 1995). In addition, dyads in winter rarely enter closed habitats (only 7% of fission events occurred in closed habitat); if caribou select closed habitats when they are either alone or in smaller groups (Webber et al., 2021), then there is little opportunity for fission events to occur in these habitats when there are fewer groups from which to split. The probability of fission and group size are two distinct aspects of grouping behavior.

Our results, in combination with past work in our system (Webber & Vander Wal, 2021), suggest that habitat openness affects group size, but not the individual probability of leaving a group.

We predicted landscape heterogeneity to induce a conflict of interest in dyads and increase the probability of fission. We used two measures of heterogeneity: composition (i.e., Shannon index: the diversity of habitat types in an area) and configuration (i.e., the contagion index: the distribution of habitat types in an area). High Shannon indices indicate landscapes with a greater diversity of land cover types, whereas a location with a higher contagion index indicates a greater number of small and disconnected patches. Landscape composition increased fission probability, while configuration had no effect, a pattern observed elsewhere (e.g. Bélisle, Desrochers, & Fortin, 2001). Taken together we submit that variable habitat types, regardless of spatial arrangement, lead to conflict of interest between group members (P2). When dyads travel through heterogeneous landscapes, the complexity of decisions about where to go next increases, thereby increasing the likelihood of disagreement between individuals regarding personal needs and motivations.

The activity budget hypothesis predicts that individuals of similar size have similar energetic requirements and more synchronous patterns of activity, which results in reduced likelihood of fission (Conradt, 1998). We did not find support for this hypothesis in our analysis, where body size difference (differences in chest girth range = 0 – 26 cm) had no effect on fission rates (P3). Furthermore, we found a contradictory pattern in social association strength for female caribou (P5), where individuals associated more closely with more differently sized conspecifics. Although we do not have relatedness or dominance hierarchy data for our population, the unexpected size-specific pattern of association we found may emerge from either kin based patterns of grouping (Djaković et al. 2012) or it could be the result of larger females associating with smaller females as a means to assert dominance (Barrette & Vandal, 1986). Indeed, caribou often form groups of loosely related kin (Djaković et al. 2012), while larger body size is often associated with dominance (Barrette & Vandal, 1986). For smaller individuals, associating with dominant individuals may provide access to higher food quality (Barrette & Vandal, 1986) via social information transfer about the location and quality of food (i.e., the conspecific attraction hypothesis: Peignier et al., 2019). This may be particularly important in the winter when snow covers lichen and lichen distribution and availability is heterogeneous (Bergerud, 1974).

As we predicted, social familiarity among females influenced dyad fission. The probability of fission decreased for dyads with stronger social preference (P4). Similarly, in domestic female sheep (*Ovis aries*), familiar individuals remain in foraging groups for longer than with unfamiliar individuals (Boissy & Dumont, 2002). Grey kangaroos (*Macropus giganteus*) also spend more time foraging with conspecifics when they are familiar rather than unfamiliar (Carter et al., 2009). Strong social bonds can result in fitness benefits. For example, social bonding enhances the life expectancy of female baboons (*Papio hamadryas ursinus*: Silk et al., 2010), and increases the reproductive success of female feral horses (Cameron, Setsaas, & Linklater, 2009). Such social bonds can also enhance anti-predatory behaviour by allowing groups to divert attention from intra-specific aggression to predator vigilance and feeding (Griffiths et al., 2004).

We examined four non-mutually exclusive ecological and behavioural factors that influence fission-fusion dynamics: perceived predation risk, habitat heterogeneity, body size, and social familiarity. Fission-fusion dynamics allow for flexibility of group sizes in animal societies, which individuals use to modulate the costs and benefits of sociality in variable environments.

Our results suggest the probability of fission increased with increasing habitat heterogeneity, while more socially familiar dyads stayed together for longer. Drivers of fission-fusion dynamics notably parallel those identified as threatening caribou population persistence. Woodland caribou are currently listed as threatened in Canada and the primary reasons for their decline are increased predation and habitat loss, which are caused by a combination of anthropogenic and natural disturbance known to fragment habitats (Festa-Bianchet et al., 2011). As a result of habitat loss, forage availability is reduced, which in turn influences caribou body condition and consequently birth rates and calf survival (Crête & Huot, 1993). Moreover, during population declines, animal social environments can change, and familiar social connections may be replaced by more ephemeral or anonymous social connections (Caro & Sherman, 2011). The effects of perceived predation risk, habitat heterogeneity, body size, and social familiarity not only have potential to affect the probability of fission, but are also among the most important causes and consequences of caribou population declines. Our work addresses the effects of these four factors on the probability of fission and falls within the mandate of the conservation behaviour framework (Berger-Tal et al., 2016); that is, to conduct behavioural research that informs conservation efforts. In a broader context, caribou conservation in Canada aims to reduce mortality (Festa-Bianchet et al., 2011). We provide evidence for how two key factors (i.e., predation and habitat heterogeneity) influence fission-fusion dynamics, a behaviour known to influence fitness outcomes in ungulates (e.g., Cameron, Setsaas, & Linklater, 2009; Vander Wal et al., 2015).

## Acknowledgements

We respectfully acknowledge the territory in which data were collected and analyzed as the ancestral homelands of the Beothuk and the Island of Newfoundland as the ancestral homelands of the Mi’kmaq and Beothuk. We thank M. Laforge, M. Bonar, and R. Huang for help in the field and members of the Wildlife Evolutionary Ecology Lab for helpful comments on previous versions of this manuscript. We also thank members of the Newfoundland and Labrador Wildlife Division, including S. Moores, B. Adams, W. Barney, and J. Neville for facilitating animal captures and for logistical support in the field. Funding for this study was provided by Natural Sciences and Engineering Research Council (NSERC) Canada Graduate Scholarship to JGH, NSERC Vanier Canada Graduate Scholarship to QMRW, and a NSERC Discovery Grant to EVW.

## Supplementary information

**Table S1:**
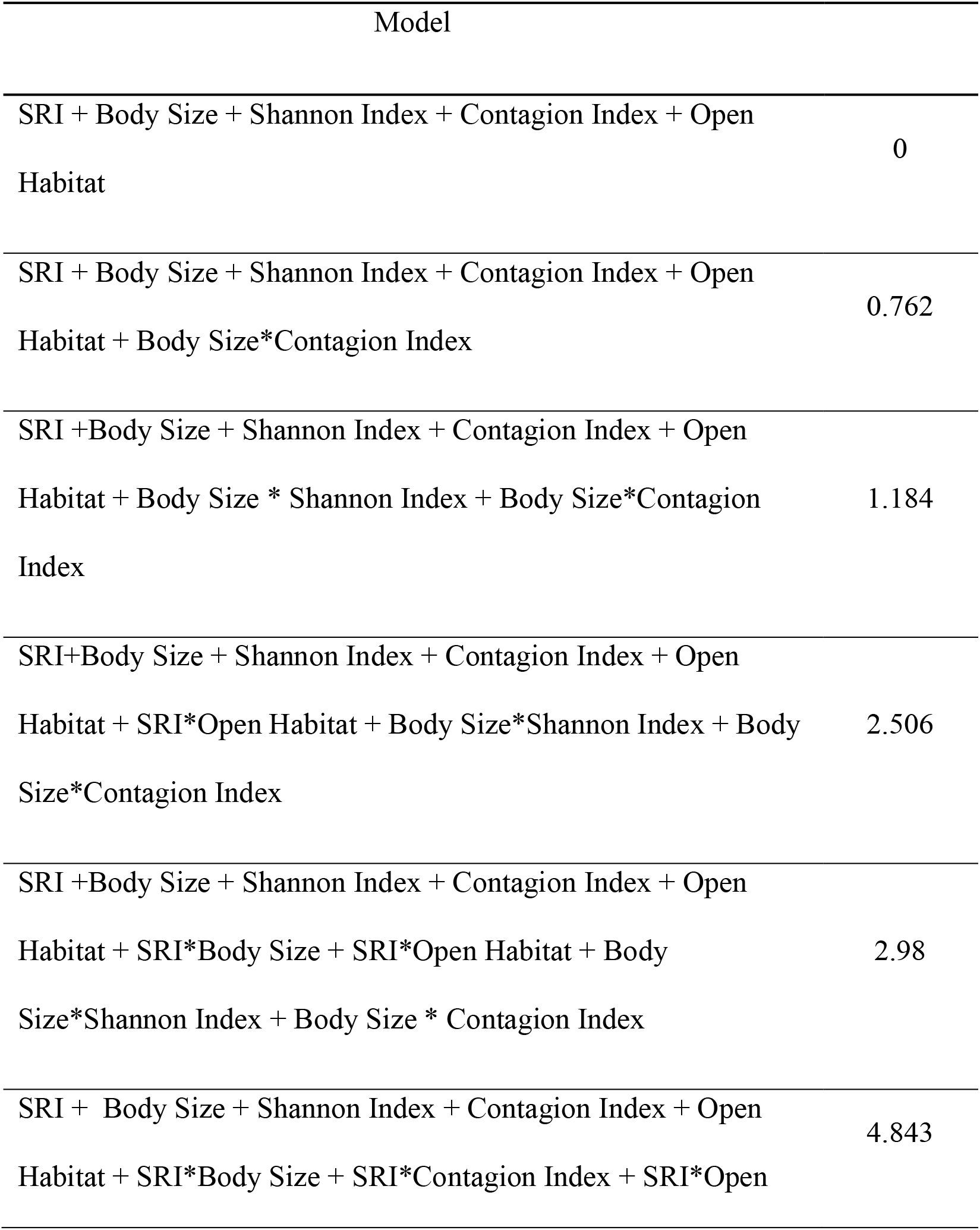
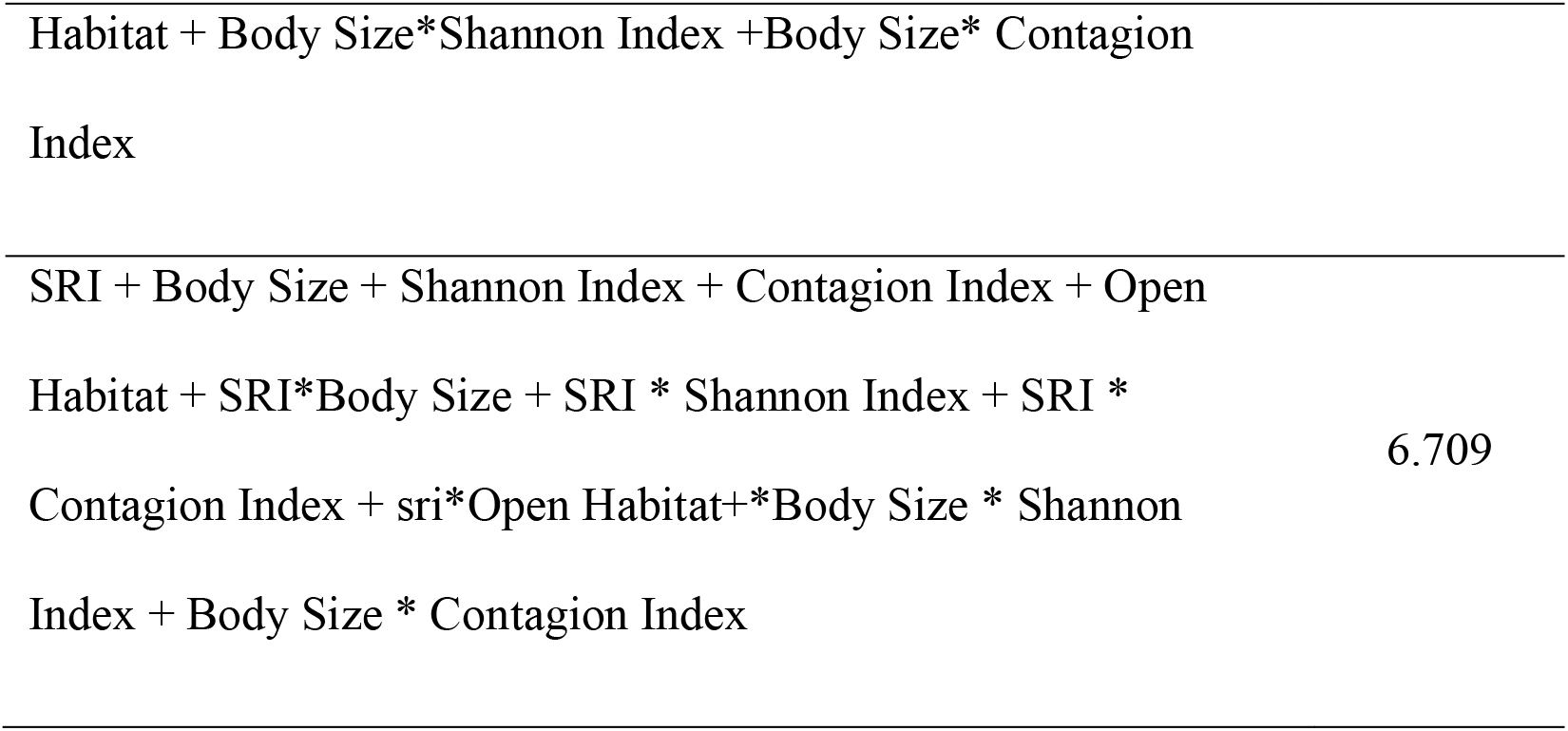
Candidate Cox proportional hazards models explaining the fission probability of caribou dyads on Fogo Island between 2017 and 2019, ranked in order of support based on AIC.

**Table S2:** Summary of our model testing the effects of home range overlap and difference in body size on the simple ratio index, that represent social familiarity of caribou in Fogo Island, Canada. Results with p < 0.05 are presented in bold.

| Simple ratio index | $\beta$ | SE | t-value | p-value |
| --- | --- | --- | --- | --- |
| Intercept | 0.096 | 0.022 | 4.333 | 0 |
| **Home range overlap** | **0.198** | **0.020** | **9.815** | **<0.0001** |
| Difference in body size | -0.003 | 0.002 | -1.446 | 0.148 |
| **Home range overlap x**<br>**Difference in body size** | **0.007** | **0.003** | **2.855** | **0.004** |
| Random variables | Variance | SD |  |  |
| Dyad ID | 0.003 | 0.062 |  |  |
| Year | 0.001 | 0.023 |  |  |
| Residual | 0.002 | 0.054 |  |  |

**Figure S1:**
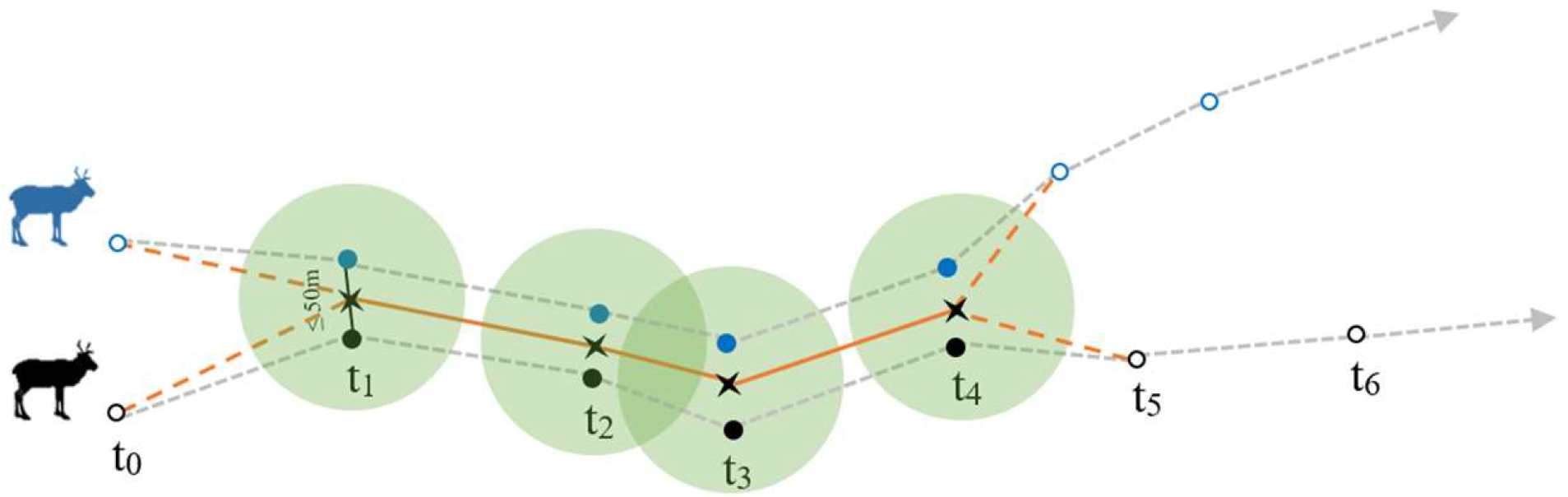
Descriptive schema of dyad fission-fusion. Black and blue points represent two different caribou moving through space and time. Each X represents the centroid of locations between the dyad and dyad steps are represented with solid orange lines. Dashed grey lines represent steps for each individual of the dyad. Our analyses of dyad space use and movement considered the shared dyad centroids and steps, not the individual paths during the dyad’s duration. Dashed orange lines represent individual paths taken by each caribou before merging in a dyad or after splitting and open circles represent caribou outside a dyad. In this schema, the dyad is created, i.e. fusion, at t_1_ because the two caribou stayed within 50m during two consecutive time-steps, t_1_ and t_2._ The dyad separates, i.e. fission, at t_4_ because the two caribou were in a dyad before t_4_ but were apart during two time-steps after, t_5_ and t_6_. Green circles represent the buffers in which time-dependent landscape metrics were calculated.

